# Genetic dissection of Rift Valley fever pathogenesis: *Rfvs2* on mouse chromosome 11 enables survival to acute-onset hepatitis

**DOI:** 10.1101/545129

**Authors:** Leandro Batista, Gregory Jouvion, Dominique Simon-Chazottes, Denis Houzelstein, Odile Burlen-Defranoux, Magali Boissière, Satoko Tokuda, Tania Zaverucha Do Valle, Ana Cumano, Marie Flamand, Xavier Montagutelli, Jean-Jacques Panthier

## Abstract

The systemic inoculation of mice with Rift Valley fever virus (RVFV) reproduces major pathological features of severe human disease, notably the acute-onset hepatitis and delayed-onset encephalitis. We previously reported that a genomic interval *(Rvfs2)* derived from the susceptible MBT/Pas strain is associated with reduced survival time after RVFV infection. In this study, we investigated the pathophysiological mechanisms by which *Rvfs2* confers increased susceptibility to BALB/c mice that are congenic for *Rvfs2* (C.MBT-*Rvfs2*) after infection with virulent RVFV. Clinical traits, biochemical parameters, and histopathological features indicated similar liver damage in BALB/c and C.MBT-*Rvfs2* mice between the third and fifth days after infection. However, C.MBT-*Rvfs2* mice died at that point from acute liver injury while most BALB/c mice recovered from this condition but eventually died of encephalitis. We observed that hepatocytes proliferated actively within the infected liver of BALB/c mice on the sixth day after infection, promoting organ regeneration on the eighth day after infection and recovery from liver damage. We found that the production of infectious virions was up to 100-fold lower in the peripheral blood and liver of BALB/c compared to C.MBT-*Rvfs2* mice. Likewise, RVFV protein amounts were much lower in BALB/c liver compared to C.MBT-*Rvfs2* liver. Primary cultured hepatocytes showed higher viral replication rate in C.MBT-*Rvfs2* which could contribute to the susceptibility conferred by *Rvfs2*. Using bone marrow chimera experiments, we uncovered that both hematopoietic and non-hematopoietic cells are required for the BALB/c allele of *Rvfs2* to exert its protective effects against the RVFV-induced acute liver disease. Taken together, we have established that *Rvfs2* acts as an important RVFV restriction factor by limiting virus multiplication in mice.

**Author Summary:** Rift Valley fever (RVF) is a mosquito-borne viral disease with potential to generate a public health emergency. The wide variation in RVF symptoms and severity observed within patient population suggests that natural host genetic determinants, among other factors, can influence the disease outcome. Infection of mice mimics several features of the pathology in humans, including acute-onset hepatitis and delayed-onset encephalitis. BALB/c inbred mice bearing the BALB/c haplotype at the *Rvfs2* locus survive longer than those bearing the MBT haplotype. In this study, we investigated clinical traits, biochemical parameters, virological evidence, and histological features to characterize the pathogenesis of RVF in early and late susceptible mice. We show that animals of both groups develop acute liver disease shortly after infection. We demonstrate that, by comparison with early susceptible mice, BALB/c mice exhibit significantly reduced replication of RVF virus *in vivo* in the blood and liver and *in vitro* in primary cultured hepatocytes, and eventually self-recover from the liver damages. We use reciprocal transplantations of bone marrow cells between early and late susceptible mice to show that survival to severe liver disease requires both hematopoietic and non-hematopoietic cells. Taken together, we establish *Rvfs2* as a single locus that enables mice to survive RVF virus-induced liver disease.

## Introduction

Rift Valley fever (RVF) is a mosquito-borne viral disease with potential to generate a public health emergency [1]. In humans, infection leads to a great variety of clinical manifestations that range from a febrile influenza-like illness to hepatitis with fatal hemorrhagic fever, encephalitis and retinitis [2, 3]. In ruminant species, a wide variation in susceptibility to RVF disease is observed among different individuals. Some infected animals suffer from unapparent or moderate febrile reactions while others develop high fevers and severe prostration, which may lead to death in the most susceptible animals [4–6]. Sequence analysis of RVF virus (RVFV) strains collected during the 1977-1979 Egyptian outbreak has shown that, although all virus isolates carried virtually identical genotypes, remarkable differences were observed in pathogenesis across human and animal populations [7]. These findings suggest that the different pathogenic phenotypes were not linked to specific mutations in the viral genome but could rather be caused by variations in dose and route of virus exposure and by host-related factors including age, sex, overall immune response, nutritional status and genetic variants.

Careful control of experimental conditions of infection in rodent models have helped establishing host genetic factors as important determinants in RVF disease severity. The infection of laboratory rodents mimics several features of RVFV-induced pathology in humans, including hepatitis with liver necrosis and meningoencephalitis [8, 9]. The first rat models consisted of the Wistar-Furth (WF) inbred strain which is highly susceptible to the hepatitis induced by subcutaneous inoculation with RVFV, while the Lewis (LEW) strain is largely resistant [8, 10]. Notably, WF rats were not uniformly susceptible to different RVFV strains [11]. Inhalation exposure to RVFV confirmed the extreme susceptibility of the WF strain to RVFV-induced hepatitis [12]. The segregation analysis of the RVFV susceptible phenotype in LEW and WF backcrosses suggested a simple Mendelian dominant control [10]. A WF.LEW congenic strain resistant to the fatal hepatitis was created by repeated backcrosses from the resistant LEW to the WF susceptible genetic background [13]. A single region on rat chromosome (Chr) 3 was shown to significantly increase the survival rate of animals carrying the LEW haplotype [14] but the gene accounting for this improved resistance has yet to be identified.

Mouse inbred strains also exhibit differences in their susceptibility to an infection with RVFV. In one study, the subcutaneous infection of BALB/c mice with 10^3^ plaque-forming units (PFU) of the ZH501 RVFV strain led to an extensive infection of the liver [9]. The resulting liver disease accounted for the death of most animals between days 3 and 6 post infection (p.i.). Mice that survived this early liver disease later developed encephalitis and died around day 8 p.i. [9]. In another study, C57BL/6 mice appeared more susceptible than BALB/c mice under similar experimental conditions and succumbed to acute liver disease within 4 days [15]. We have tested the susceptibility of additional strains derived from various *Mus musculus* subspecies trapped in the wild. The most severely affected strain within this collection, MBT/Pas (MBT), developed very early onset RVF disease. After intraperitoneal infection with 10^2^ PFU of virulent RVFV strain, either Egyptian ZH548 or Kenya 98, MBT mice died more rapidly than BALB/c or C57BL/6 mice [16]. It is worth noting that MBT mice are susceptible to RVFV but resistant to several other viruses [16], suggesting that the susceptibility to RVFV exhibited by MBT mice is not attributable to generalized immunodeficiency. In flow cytometry studies, we have recently shown that MBT mice displayed several immunological alterations after RVFV infection. Furthermore, these mice failed to prevent high viremia and viral antigen loads in the blood, spleen, and liver [17]. We also showed that, in MBT mice, RVF susceptibility is inherited in a complex polygenic fashion and we identified three genomic intervals on Chr 2, 11 and 5 affecting survival time after RVFV infection. Each of these MBT-derived intervals, designated Rift Valley fever susceptibility 1 (*Rvfs1*), *Rvfs2* and *Rvfs3* respectively, conferred reduced survival time in C.MBT congenic strains in which these intervals had been transferred onto the less susceptible BALB/c genetic background [18]. The pathogenic mechanisms for the early death induced by RVFV in the C.MBT congenic strains are currently unknown.

In this study, we investigated the phenotypic features associated with morbidity in BALB/c mice congenic for the MBT-derived *Rvfs2* interval, i.e. C.MBT-*Rvfs2* mice. We focused our investigations on male mice which exhibit slightly higher susceptibility to RVFV infection [18]. The study of clinical, biochemical and virological parameters, as well as histopathological features of the RVF disease showed that mice from both BALB/c and C.MBT-*Rvfs2* inbred strains exhibited hepatic disease. The first clinical signs of disease were detected on the third day of infection in both strains. However, while C.MBT-*Rvfs2* mice began to die on day 4 of acute liver disease, most BALB/c mice recovered and died three to nine days later of encephalitis. Since MBT-derived *Rvfs2* allele was associated with increased viral load in the liver and higher viral replication rate in primary cultured hepatocytes, we conclude that the death of C.MBT-*Rvfs2* mice is due to enhanced susceptibility to the acute-onset, RVFV-induced liver disease. Reciprocal transplantations of bone marrow cells between BALB/c and C.MBT-*Rvfs2* mice showed that both hematopoietic and non-hematopoietic cells are required for the capacity of BALB/c mice to survive liver damages.

## Results

### BALB/c and C.MBT-*Rvfs2* mice have evidence of liver disease at the early stage of infection

C.MBT-*Rvfs2* congenic mice carry a ≈17 Mb segment of chromosome 11 region from the MBT strain on the BALB/c background (Fig 1A) [18]. The challenge of C.MBT-*Rvfs2* mice with 10^2^ PFU of the ZH548 RVFV strain showed that *Rvfs2* has a strong effect on susceptibility to RVFV. More than 50% of C.MBT-*Rvfs2* mice died within 4 days after infection with RVFV, whereas half of BALB/c mice survived for over 8 days (Mantel-Cox’s Logrank test, P<0.0001) (Fig 1B).

**Figure 1.**
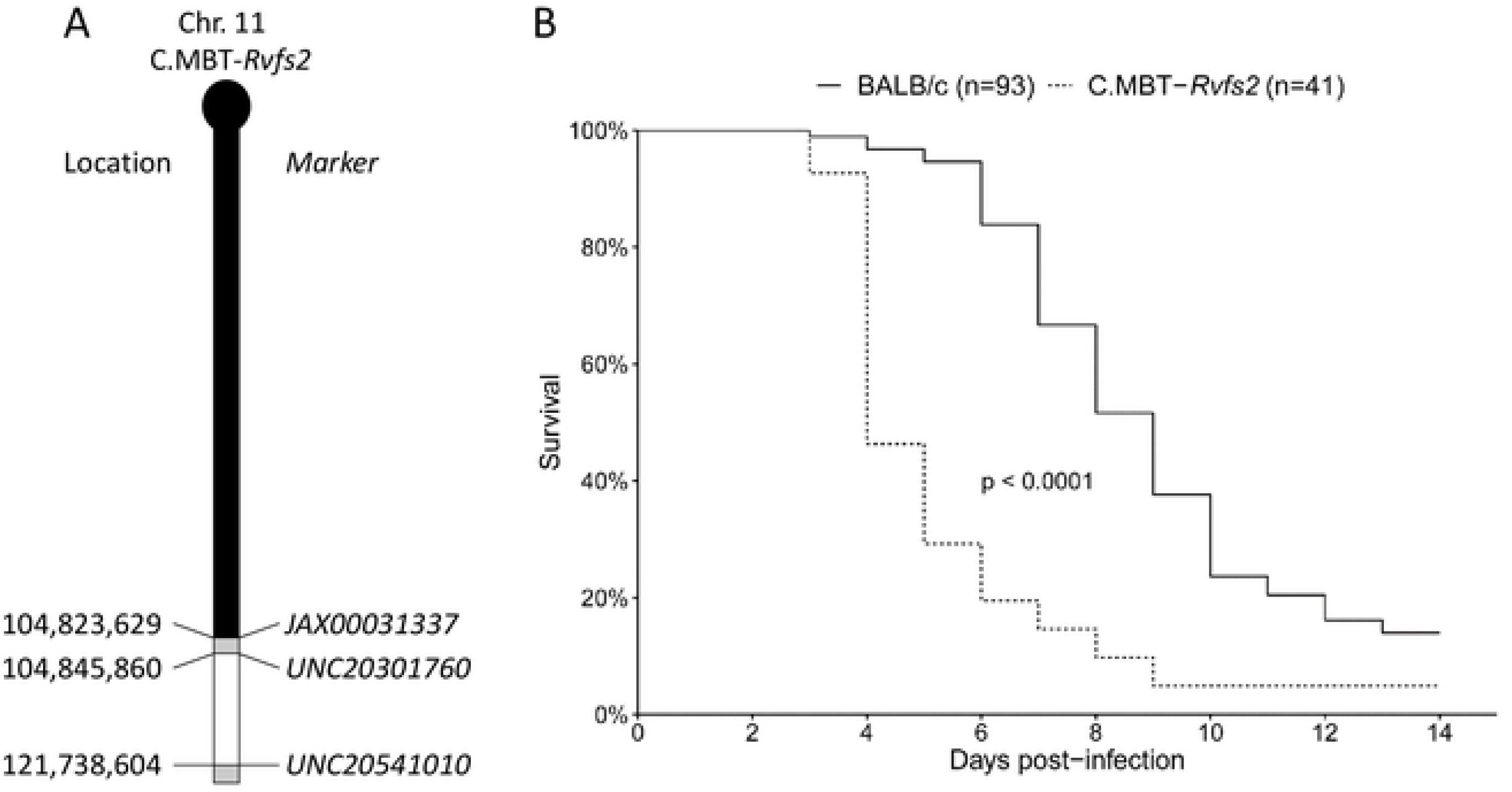
Representation of the MBT-derived *Rvfs2* region in the congenic C.MBT-*Rvfs2* strain and its effect on mouse survival. (A) Haplotype structure of the congenic segment of chromosome 11 in C.MBT-*Rvfs2* (Rvfs2) strain. The MBT-derived segment is depicted in white on the BALB/c chromosome 11 background (black). Regions of unknown genotype are depicted in grey. Markers are SNPs from the GigaMUGA array (https://support.illumina.com/downloads/geneseek-ggp-giga-muga-product-files.html) and position are given in bp from mouse Genome Build 37 (corrected from [18]). (B) Survival curves of C.MBT-*Rvfs2* and BALB/c male mice infected with 100 pfu IP (Mantel-Cox’s Logrank test; p<0.0001).

Clinical signs of disease in RVFV-infected C.MBT-*Rvfs2* mice were monitored daily in comparison with BALB/c in order to explore the causes of this increased susceptibility. C.MBT-*Rvfs2* mice developed signs of an acute disease with ruffled fur, hunched appearance and lethargy as early as days 3 and 4 p.i. In contrast, most RVFV-infected BALB/c mice exhibited the first symptoms of disease later, from day 6 p.i. BALB/c mice showed different degrees of disorders, including ascending paralysis, ataxia, head-tilt and circling behavior. These clinical observations suggest that distinct presentations of the disease are controlled be the introgressed chromosomal region.

Quantitative clinical traits and biochemical parameters associated with the health status were recorded to further characterize differences in disease progression between BALB/c and C.MBT-*Rvfs2* mice. Body weight and body temperature were measured daily in RVFV-infected mice from both strains. On average, BALB/c mice lost body weight on days 3 and 4, and from day 7 to day 9 p.i. Between these two intervals their body weight remained stable. By contrast, C.MBT-*Rvfs2* mice lost weight rapidly from day 3 p.i. until their death (Fig 2A). No differences were found between the two strains in body weight loss at days 3 and 4 (two way ANOVA, P(strain effect)=0.54). Temperature measurements indicated that neither BALB/c nor C.MBT-*Rvfs2* mice became febrile during the course of infection. A significant drop in body temperature was observed one day before death in both inbred strains, regardless of the cause of death (Fig 2B). Overall, no differences in body temperature between BALB/c and congenic mice were found (two-way ANOVA, P(strain effect)=0.69). As RVFV is known to be a hepatotropic virus [19], blood levels of liver enzymes were measured during the disease course in uninfected and infected BALB/c and C.MBT-*Rvfs2* mice. Alanine aminotransferase (ALT) and aspartate transaminase (AST) peaked on day 4 p.i. in both infected strains (Fig 2C and 2D), indicating hepatocyte damage. After day 5 p.i., AST and ALT serum levels decreased slowly in infected BALB/c mice and returned to normal levels on day 8 p.i., suggesting recovery from the liver disease. Altogether the development of the RVF disease in the first 4 days was similar in both inbred strains.

**Figure 2.**
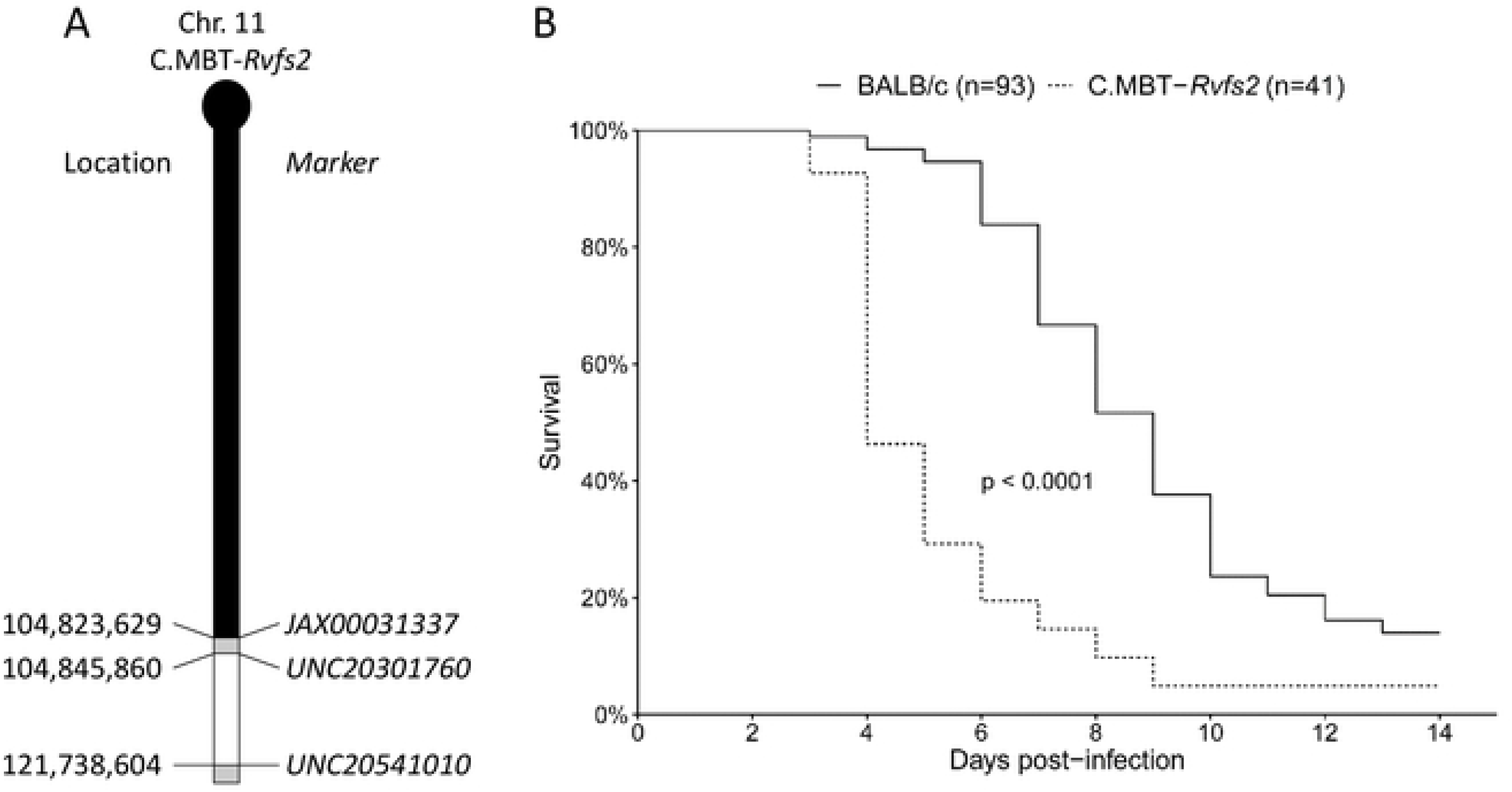
Clinical traits and biochemical parameters of RVFV-infected C.MBT-*Rvfs2* and BALB/c mice. (A) Daily body weight variation in C.MBT-*Rvfs2* (Rvfs2) and BALB/c mice after infection with RVFV ZH548 (mean ± SEM). Positive values indicate weight gain (in %) from previous day while negative values indicate weight loss. (B) Body temperature variation in C.MBT-*Rvfs2* and BALB/c mice during the days preceding the death (mean ± SEM). No difference was observed between the two strains (two-way ANOVA, p=0.69). (C-D) Alanine aminotransferase (ALT) (C), and aspartate aminotransferase (AST) (D) levels in the serum of C.MBT-*Rvfs2* and BALB/c mice. By day 5 p.i., all C.MBT-*Rvfs2* mice had died. Data are means ± SEM for N= 4-9 mice per day, except for day 8 in BALB/c mice where N=1.

### BALB/c and C.MBT-*Rvfs2* mice exhibit liver damage at days 3 and 4 p.i

We studied in further detail the tissue damage caused by RVFV in the liver of infected BALB/c (N=10) and C.MBT-*Rvfs2* (N=8) mice euthanized on day 3 p.i. Histopathological analyses of the liver revealed three different lesion profiles of increasing severity in both mouse genotypes (Fig 3). Five out of 10 BALB/c and 5/8 C.MBT-*Rvfs2* mice exhibited mild, multifocal and well demarcated lesions defined as Profile 1. Liver lesions in these mice were characterized by hepatocyte cell death associated with small inflammatory infiltrates containing fragmented neutrophils (Fig 3A and 3B). Immunohistochemical (IHC) labeling directed against the viral N protein was used to identify infected cells (note that the technique used could not provide quantitative information on the infection level in infected cells). This analysis revealed small multifocal foci (less than 100 μm in diameter) of infected hepatocytes (Fig 3C). Profile 2 was observed in 3/10 BALB/c and 2/8 C.MBT-*Rvfs2* mice. This profile was also characterized by multifocal lesions with hepatocyte cell death. However, lesions were more severe, extensive and diffuse (Fig 3D and 3E). IHC analyses detected a stronger signal with slightly larger foci of infected hepatocytes (Fig 3F). A third profile was observed in 2/10 BALB/c and 1/8 C.MBT-Rv/ŝ2 mice. Liver sections categorized as Profile 3 displayed severe and diffuse tissue damage without signs of inflammation. Lesions were characterized by acute and massive cell death of hepatocytes, numerous viral inclusion bodies in the nuclei of cells (Fig 3G and 3H), and a diffuse positive immunolabeling of hepatocytes for the viral N protein (Fig 3I). None of these lesions were observed in liver sections of uninfected BALB/c and C.MBT-*Rvfs2* mice. Collectively, these results indicated that BALB/c and C.MBT-*Rvfs2* mice experienced similar liver conditions on day 3 p.i. with the same range of histological lesions, from mild to severe, up to extensive destruction of the liver parenchyma. Overall, non-quantitative IHC indicated, in each profile, similar amounts of RVFV-infected liver cells in both strains.

**Figure 3.**
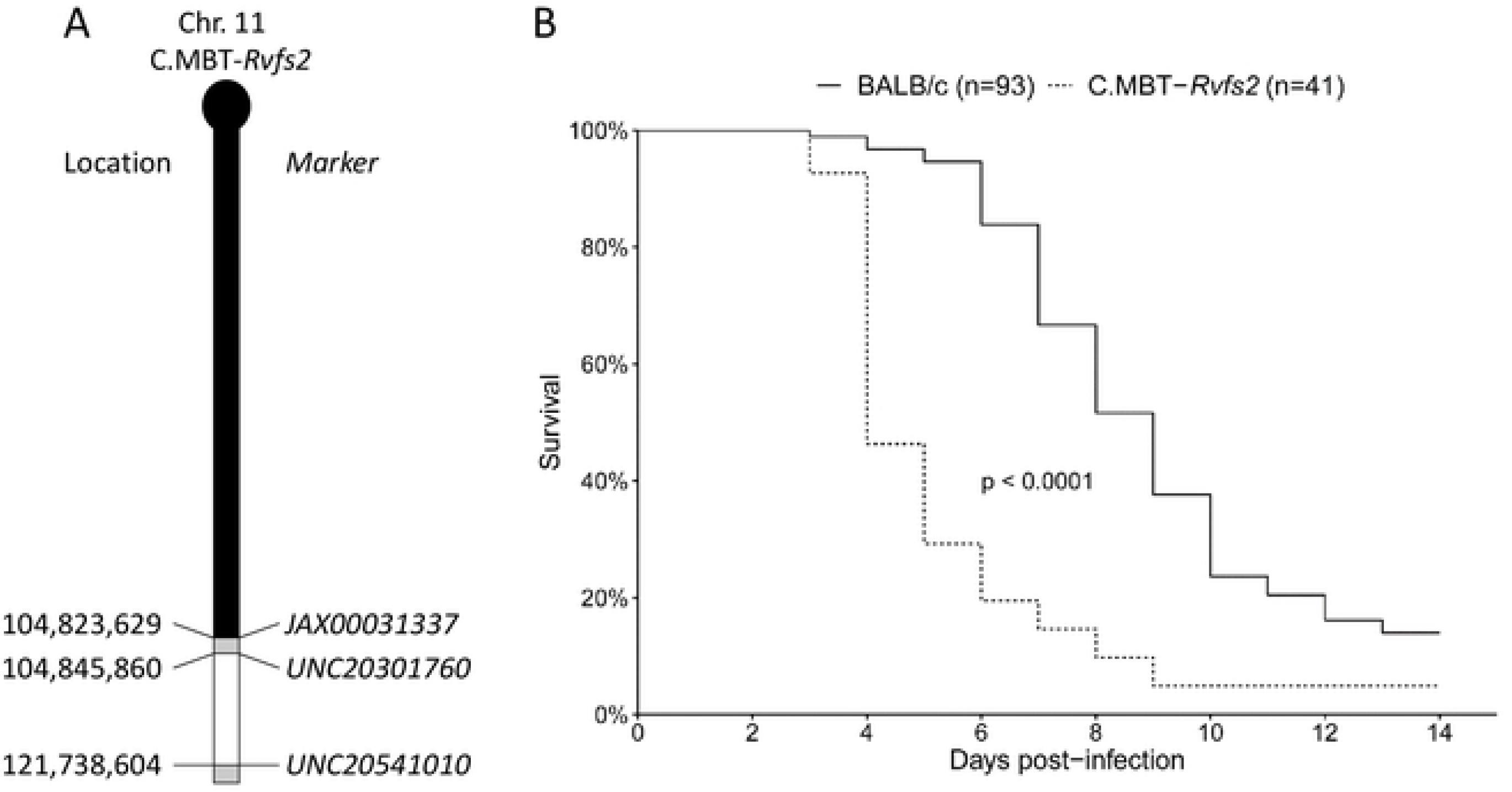
Histopathology and immunohistochemistry analyses of liver from BALB/c and C.MBT-*Rvfs2* mice on day 3 p.i. Three distinct histological profiles were found in 10 BALB/c and 8 C.MBT-*Rvfs2* infected mice. Profile 1: (A) Randomly distributed, multifocal inflammatory lesions (arrowheads) with (B) small well-delimited foci of necrotic/apoptotic hepatocytes associated with neutrophil infiltration. (C) Small clusters of RVFV N protein-positive hepatocytes recognized by immunohistochemistry. Profile 2: (D) Multifocal inflammatory lesions randomly distributed in the liver (arrowheads) with (E) more extensive and severe foci of necrotic/apoptotic hepatocytes than in Profile 1, without inflammatory infiltration. (F) Slightly larger clusters of N-positive hepatocytes observed after immunohistochemistry staining. Profile 3: (G, H) Massive necrosis/apoptosis of hepatocytes, (I) with a strong and diffuse immunohistochemistry staining for RVFV N protein in the parenchyma. None of these lesions were observed in the liver of uninfected BALB/c and C.MBT-*Rvfs2* mice. A, B, D, E, G, H: Hematoxylin and eosin staining; C, F, I: Immunohistochemistry for RVFV N protein.

We then examined the liver of moribund C.MBT-*Rvfs2* mice in order to provide further histological evidence of disease progression. Five congenic mice were euthanized when exhibiting clinical signs of severe disease, on days 3 (N=1), 4 (N=2) and 4.5 (N=2) p.i. All five livers displayed severe and non-inflammatory lesions, characterized by massive and acute cell death. These observations resembled closely those described above as Profile 3 (not shown), supporting the hypothesis that, in C.MBT-*Rvfs2* mice, the disease progressed from mild inflammation to non-inflammatory liver lesions with extensive tissue damage. Lesions observed in the liver of moribund C.MBT-*Rvfs2* mice were sufficient to alter liver function, leading to the rapid death of infected animals.

### Infected BALB/c mice survive the early-onset liver disease, but succumb later to encephalitis

The gradual decrease of liver transaminases between days 5 and 8 post-infection in BALB/c mice suggested hepatic tissue regeneration. We further evaluated the extent of liver recovery, by examining histopathological changes in livers of moribund BALB/c mice (N=4) euthanized between day 6 and 9 p.i. (Figs 4 and 5). On day 6 p.i., only minimal lesions were observed (Fig 4A and 4B) and few RVFV N-positive hepatocytes were detected (Fig 4C). An increased number of mitosis as well as a strong and diffuse expression of Ki67 confirmed the proliferation of hepatocytes (Fig 4D). By day 8, minimal to mild, subacute to chronic inflammatory lesions were scattered in the liver parenchyma or centered on portal tracts and consisted of small infiltrates of lymphocytes, plasma cells and macrophages (Fig 5A and 5B). Very few hepatocytes were labeled positively for the RVFV N protein, confirming an efficient viral clearance in the hepatic tissue (Fig 5C). Since BALB/c mice exhibited clinical neurological signs, we investigated their brain for infection-related lesions. Histopathological lesions were visible in the brain of moribund BALB/c mice. The virus targeted different brain anatomic structures in each individual mouse, and no pathognomonic lesion profile could be defined (Fig 5D-I). We detected (i) subacute leptomeningitis with multifocal infiltration of the leptomeninges by lymphocytes, plasma cells, and neutrophils (Fig 5D), and (ii) cell death foci in different locations of the cerebral grey matter, e.g. the outer granular layer or different brain nuclei (Fig 5E, 5F, and 5G). In these foci, shrinkage of neurons, gliosis, infiltration of neutrophils and strong RVFV N protein immunolabeling of neurons were observed (Fig 5H and 5I). These lesions were likely the cause of the neurological symptoms and eventual death in BALB/c mice.

**Figure 4.**
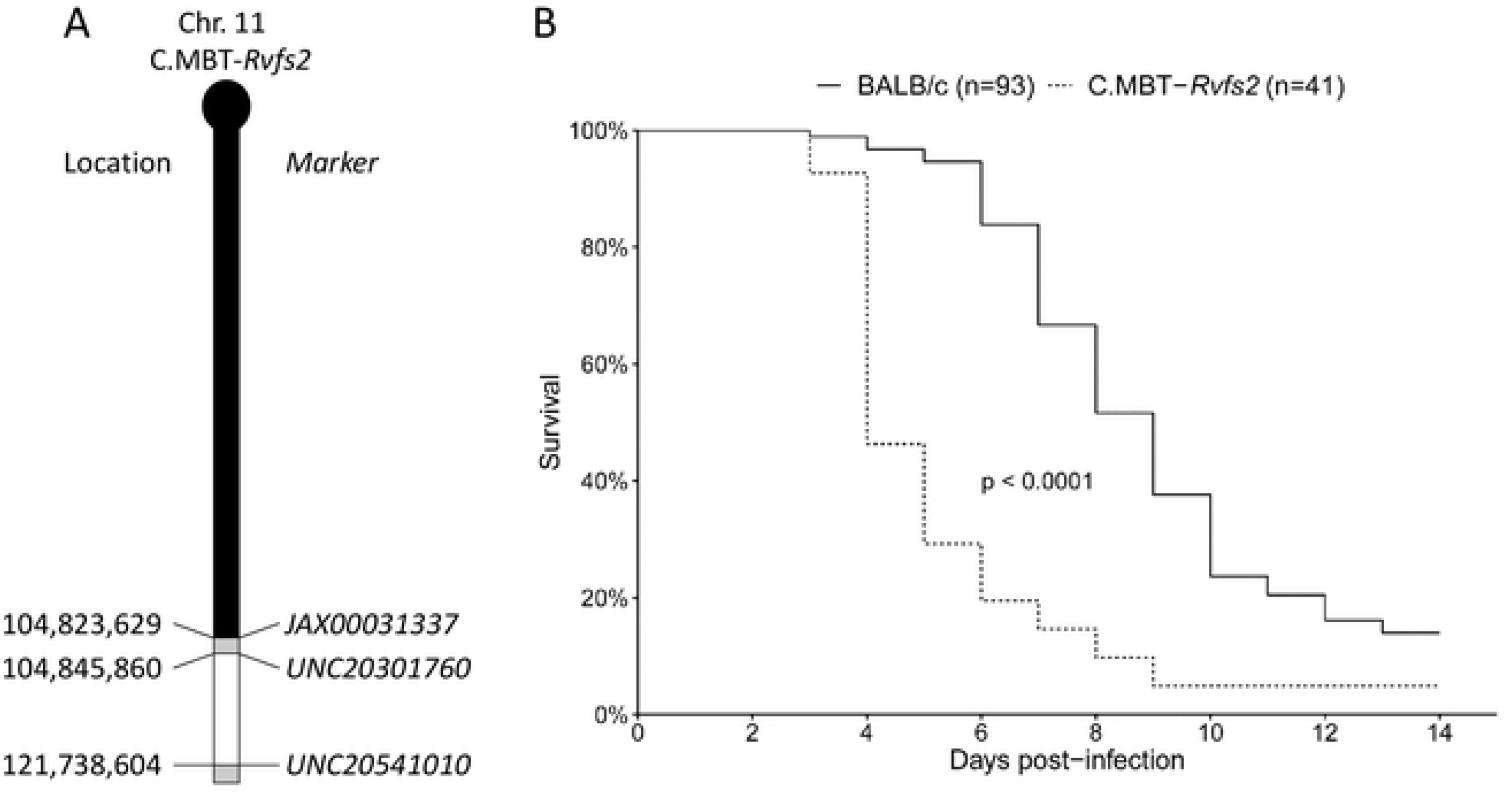
Hepatocyte proliferation and liver regeneration in BALB/c mice recovering from RVFV-induced liver disease. Liver sections of four BALB/c mice were examined at day 6 post-infection. (A) Rare and randomly distributed lesions in the liver parenchyma are observed (arrows). (B) Small infiltrates of inflammatory cells (primarily neutrophils) associated with focal hepatocyte destruction may be observed in the lesions. Increased mitotic activity is seen among hepatocytes (arrowheads). (C) Immunohistochemistry for RVFV N protein reveals a weak signal, only detected in the small foci identified in hematoxylin and eosin-stained sections (black circles). (D) Immunohistochemistry for Ki67 highlights a marked, diffuse proliferation of the hepatocytes (arrowheads). A, B: Hematoxylin and eosin staining; C: Immunohistochemistry for RVFV N protein; D: Immunohistochemistry for Ki67.

**Figure 5.**
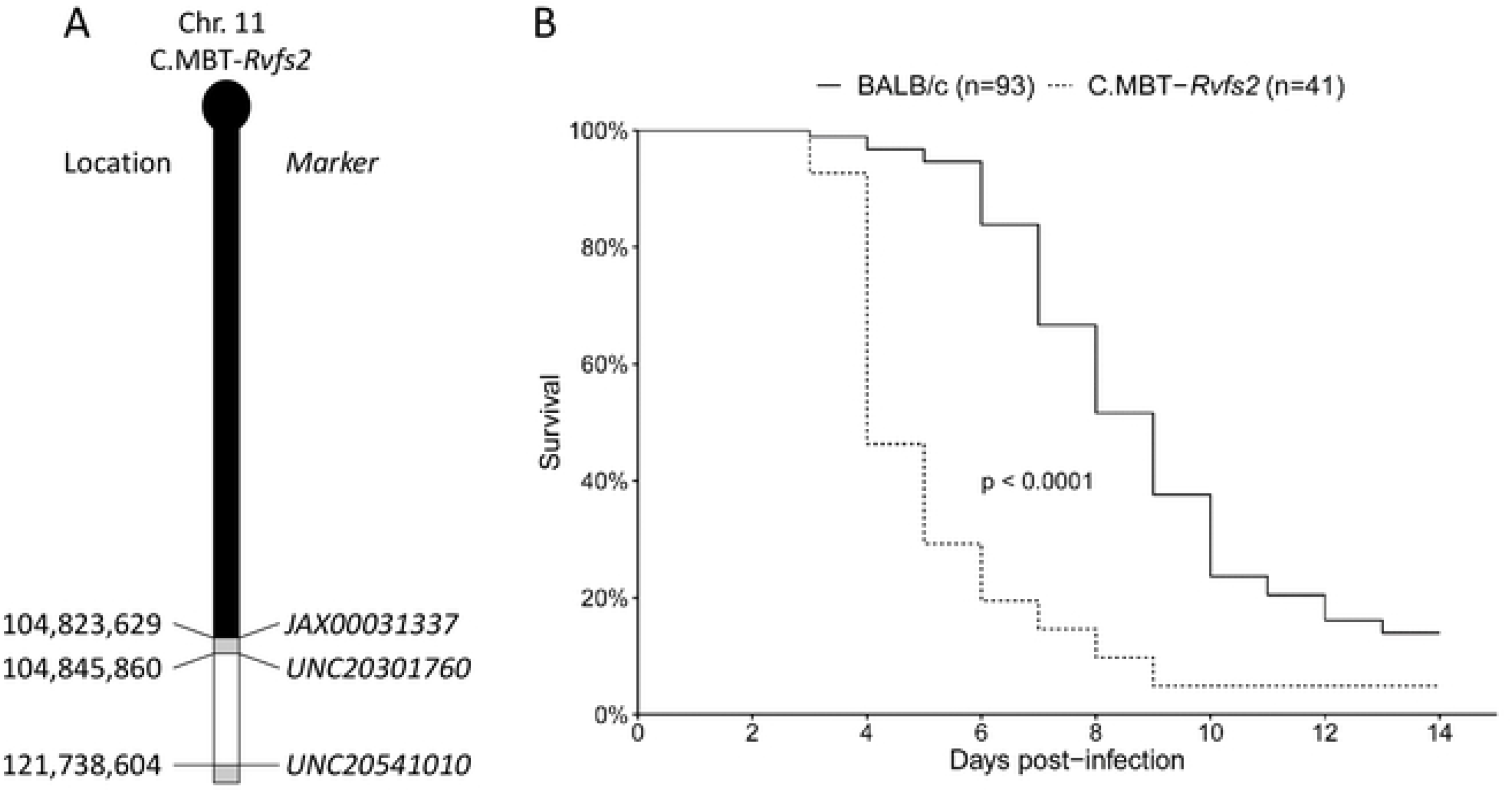
Histopathology and immunohistochemistry analyses of liver and brain from moribund BALB/c. (A-C) Liver from a moribund BALB/c mouse on day 8.5 p.i. displays minimal multifocal inflammatory lesions either randomly distributed in the liver parenchyma (arrowhead) (A) or centered on portal tracts, mostly around bile ducts (arrowhead) (B). Rare RVFV-infected cells are indicated by IHC with antibodies against RVFV N protein (C). (D-I) Brains from moribund BALB/c mice on days 7 to 9 p.i. display different inflammatory and apoptotic/necrotic lesions: subacute leptomeningitis characterized by infiltration of leptomeninges by lymphocytes, plasma cells and neutrophils (D), laminar apoptosis/necrosis of neurons in the cortical outer granular layer (E) with RVFV N protein-positive neurons (F), necrotic/apoptotic foci in different locations of the cerebral grey matter with gliosis (G) and infiltration of neutrophils (inset), and strong signal for RVFV N protein (H-I). Histology and immunohistochemistry results shown are representative of experiments performed on at least 4 animals. A, B, D, E, G: Hematoxylin and eosin staining; C, F, H, I: Immunohistochemistry for RVFV N protein.

### Elevated viral burden in the blood and liver of C.MBT-*Rvfs2* mice

In order to assess differences in the viral production, we first measured the titer of infectious viral particles in the blood and liver of BALB/c and C.MBT-*Rvfs2* mice on day 3 p.i. by standard plaque assay. The viral titers were about 80-and 100-fold higher in the C.MBT-*Rvfs2* blood and liver, respectively, compared with those found in BALB/c mice (Mann Whitney-U test, P<0.001 and P=0.016, respectively; Fig 6A and 6B). Finally, semi-quantitative protein analysis of liver extracts at day 3 p.i. by Western blot indicated that high levels of N nucleocapsid and NSs nonstructural viral proteins were found in the liver of C.MBT-*Rvfs2*, while both viral proteins were undetectable in BALB/c liver (Fig 6C) despite the same proportions of RVFV-infected cells in the two strains revealed by IHC (Fig 3). Altogether, these results indicated that, compared to BALB/c mice, C.MBT-*Rvfs2* mice have reduced ability to limit the replication of RVFV thus allowing the production of the virus systemically and specifically in the liver at much higher levels.

**Figure 6.**
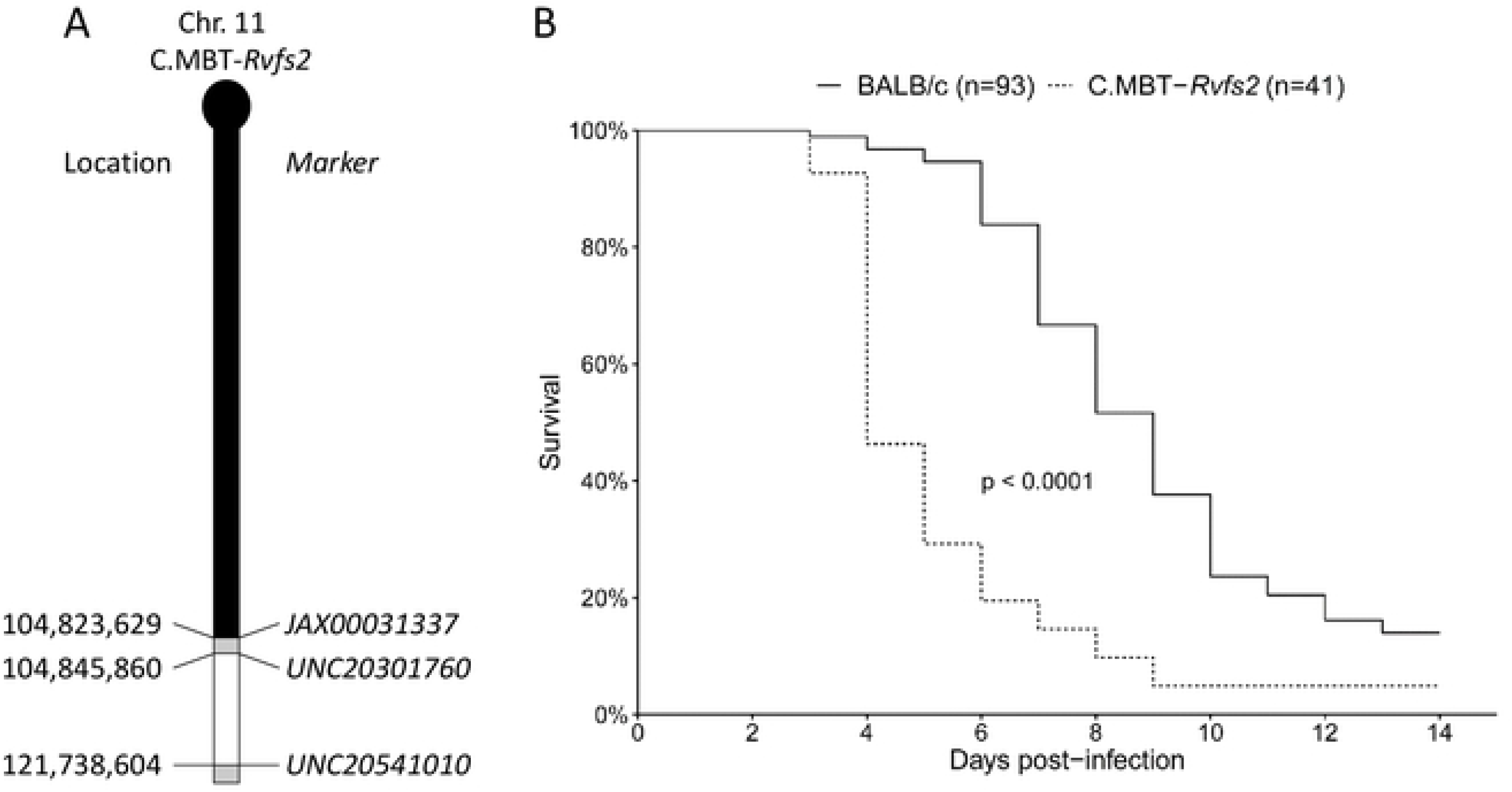
Production of viral particles and viral proteins in BALB/c and C.MBT-*Rvfs2* mice on day 3 p.i. (A) Viremia in RVFV-infected C.MBT-*Rvfs2* (Rvfs2) (N=10) and BALB/c (N=12) mice. (B) Viral titers in liver from C.MBT-*Rvfs2* (N=9) and BALB/c (N=13) mice (C) Western blotting analysis of the liver from BALB/c and C.MBT-*Rvfs2* (Rvfs2) uninfected (N=1) and infected mice on day 3 p.i. (N=2, from the groups of mice analyzed in A and B and identified as b1, b2 for BALB/c and r1, r2 for Rvfs2). RVFV-infected AML12 cells were included as a positive control. Proteins were analyzed with antibodies against NSs and N viral proteins, and betaactin. The molecular weight and positions of the marker bands (middle lane), and NSs, N and β-actin proteins (right lane) are indicated.

### Increased viral replication in C.MBT-*Rvfs2*-derived cultured primary hepatocytes

To further analyze the increased susceptibility of C.MBT-*Rvfs2* mice to RVFV-induced liver disease, we derived primary cultured hepatocytes from the liver of BALB/c and C.MBT-*Rvfs2* uninfected mice. We measured the kinetics of viral production in the culture medium over 60 h after infecting hepatocytes with RVFV. While the viral titer remained constant at 300-400 PFU/ml in the BALB/c culture, it peaked in C.MBT-*Rvfs2*-derived hepatocytes at almost 900 PFU/ml 24h after infection before decreasing at 48 and 60 hours post-infection (Fig 7). This increased viral replication in C.MBT-*Rvfs2* hepatocytes is likely one of the mechanisms responsible for the enhanced susceptibility to the liver disease conferred by the *Rvfs2* locus.

**Figure 7.**
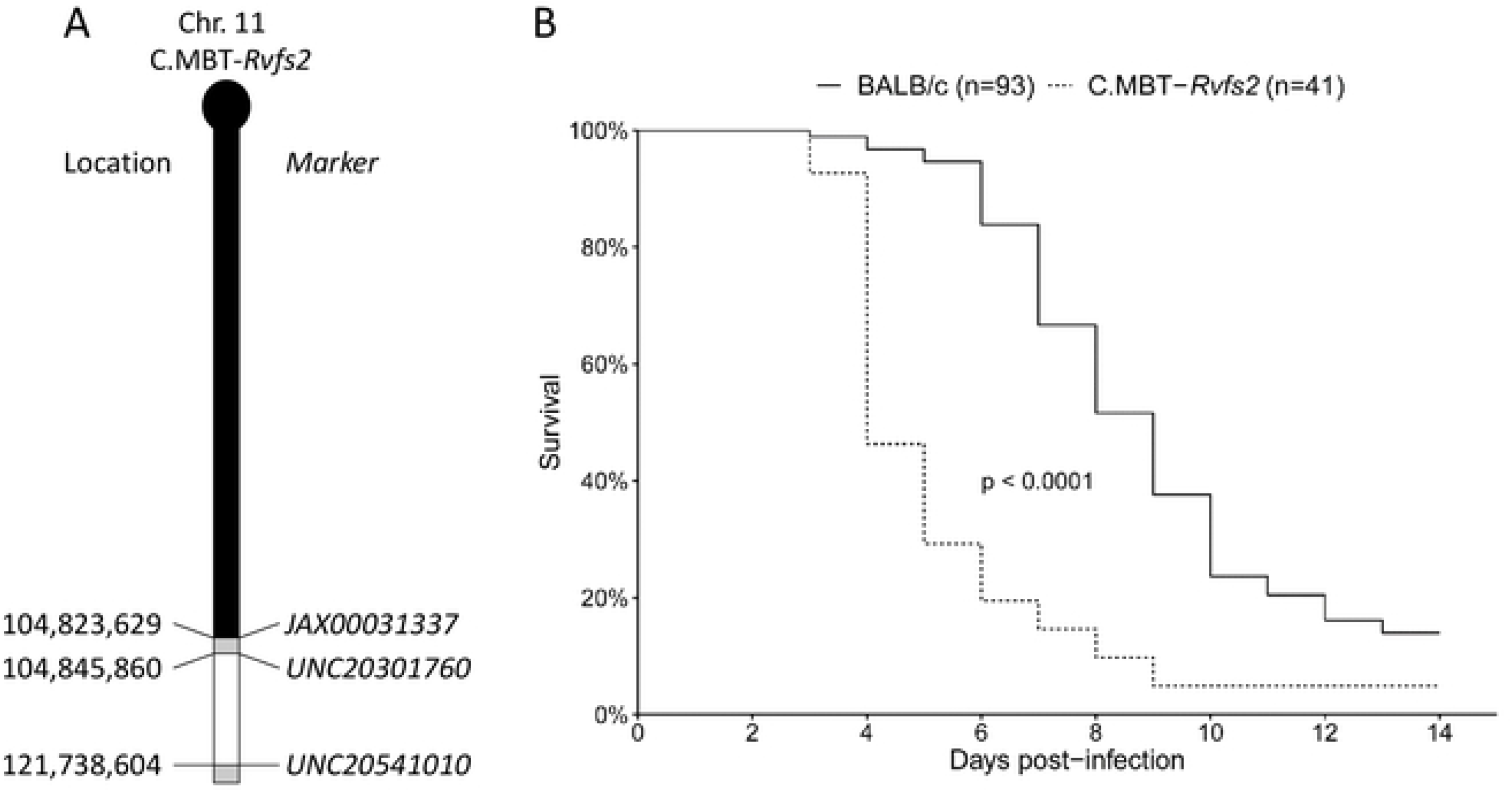
Viral replication in primary cultured hepatocytes from BALB/c and C.MBT-*Rv/f2* mice. Virus titer measured in the supernatant of primary cultured hepatocytes, at 15, 24, 48 and 60 hr p.i. with RVFV at MOI of 3. Virus titer was significantly higher in C.MBT-*Rvfs2* than in BALB/c hepatocytes at 24 and 48 hr.

### Both hematopoietic and non-hematopoietic cells are required for *Rvfs2*-dependent survival to liver disease

We have recently shown that the inbred strain MBT displays multiple immune-related defects in response to RVFV infection [17]. Together with transcriptomics data from a previous study [16], these results suggest that innate immune cells might be the critical determinant for survival to liver disease. To test this hypothesis, we produced chimeric mice using crosswise transplantations of bone-marrow cells after total body irradiation to evaluate whether the effects of *Rvfs2* in hematopoietic cells, in non-hematopoietic cells, or in both were required for survival to the RVFV-induced liver disease. We generated BALB/c mice reconstituted with C.MBT-*Rvfs2* marrow (C.MBT-*Rvfs2* → BALB/c chimeras) and C.MBT-*Rvfs2* mice reconstituted with BALB/c marrow (BALB/c → C.MBT-*Rvfs2* chimeras). Controls consisted of irradiated mice reconstituted with isogenic marrow, (BALB/c → BALB/c and C.MBT-*Rvfs2* → C.MBT-*Rvfs2* chimeras). After reconstitution, the mice were infected with RVFV. As shown in Figure 8, BALB/c → BALB/c mice survived significantly longer than C.MBT-*Rvfs2* → C.MBT-*Rvfs2* mice (Mantel-Cox’s Logrank test, P=0.0002), like the nonmanipulated BALB/c and C.MBT-*Rvfs2* strains (Fig 1B). Interestingly, the survival time was significantly shorter in C.MBT-*Rvfs2* → BALB/c mice compared to BALB/c → BALB/c mice (Mantel-Cox’s Logrank test, P<0.01), suggesting that bone marrow-derived cells are needed to confer prolonged survival in BALB/c mice. However, the survival time of BALB/c → C.MBT-*Rvfs2* mice was not increased compared to C.MBT-*Rvfs2* → C.MBT-*Rvfs2* mice (Mantel-Cox’s Logrank test, P=0.78), suggesting that bone marrow-derived cells from BALB/c mice alone are not sufficient to confer the BALB/c phenotype to C.MBT-*Rvfs2* mice. Together, these results suggest that the *Rvfs2* effects in both hematopoietic and non-hematopoietic compartments are critical for survival to the early-onset liver disease in RVFV-infected mice.

**Figure 8.**
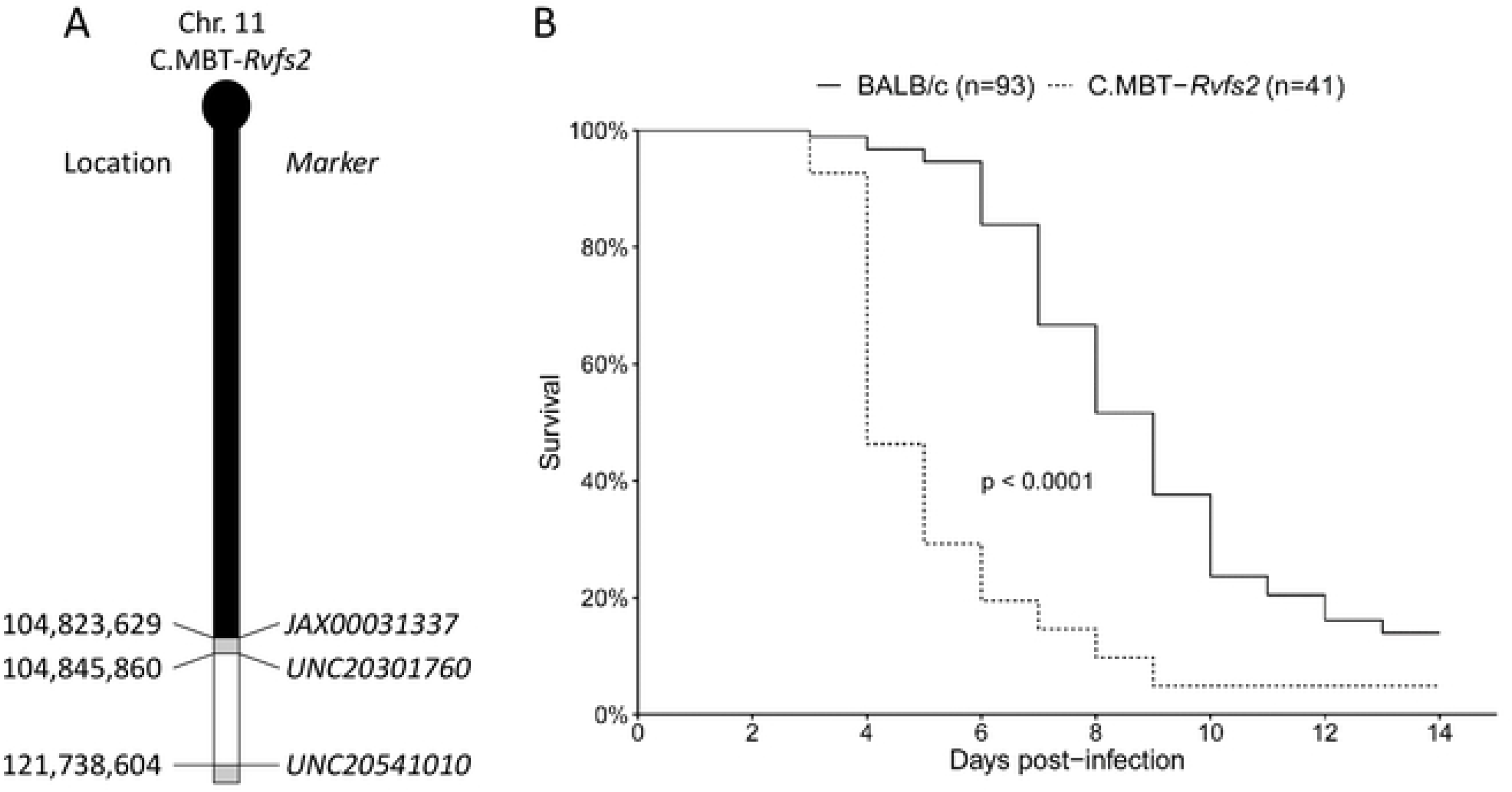
Survival curves of chimeric mice generated by reciprocal transplantation of bone marrow cells. Sub-lethally irradiated C.MBT-*Rvfs2* (*Rvfs2*, red lines) or BALB/c (black lines) recipient mice received ~ 3×10^6^ bone marrow cells from either C.MBT-*Rvfs2* (dashed lines) or BALB/c (solid lines) donor mice on the same day as irradiation. The recipient mice were infected intraperitoneally with 10^2^ PFU RVFV six weeks later. Asterisks refer to the comparison between each group and the BALB/c → BALB/c control group (Mandel-Cox’s Logrank test; **P<0.01, ***P<0.001).

## Discussion

There is considerable variability in the ability of patients and livestock to survive RVF disease. A number of factors, such as the viral strain, the route and dose of viral exposure and the age, sex, nutritional and immune status of the host, can modulate the severity of the disease and contribute to the balance between recovery and death, a complex phenotype that involves multiple systems, organs, tissues, immune cells and cellular pathways, under the influence of multiple genes. The importance of inoculation route and dose and of the age and sex of the host has been experimentally demonstrated in laboratory rodents [10, 12, 13, 16, 20]. Experiments in laboratory rodents have further demonstrated that survival time and survival rate following RVFV infection are influenced by host genetic determinants [10, 16]. Our previous studies have shown that susceptibility to RVF in the inbred MBT mouse strain is inherited in a complex fashion, with sex influencing the severity of the disease. Three RVF susceptibility loci *(Rvfs)* with a moderate effect on the survival time were identified [18]. The introgression by repeated backcrosses of each of these chromosomal regions within the less susceptible genetic background BALB/c led to congenic mice exhibiting significant reduction in survival time compared to the BALB/c control groups [18]. Functional studies are needed to unravel the nature and the role of the genes within the *Rvfs* loci in the pathogenesis of RFV. We chose to focus our efforts first on *Rvfs2,* a 17Mb genomic interval, because of its strongest effect.

In our mouse model, the animals are infected by intraperitoneal injection of 10^2^ PFU of the ZH548 RVFV strain [2, 21]. This virus dose was initially chosen to induce high mortality in both MBT and BALB/c parental strains (S1 Table). In these conditions, most MBT mice died within 5 days p.i. with the clinical signs of liver disease. By contrast, most BALB/c mice lived beyond that date and exhibited signs of encephalitis, such as paralysis, ataxia, or head-tilting behavior [16]. Notably, the difference in the days of death between BALB/c and MBT mice was identical at infectious doses ranging from 10 to 1000 PFU (S1 Table).

The pathogenesis induced by the subcutaneous challenge of BALB/c mice with 10^3^ PFU ZH501 RVFV has been recently characterized in detail [9, 22]. Approximately 80% of BALB/c mice infected in these conditions were reported to have succumbed with severe liver disease between days 3 and 6 p.i., a much higher percentage than the one observed in our study (<10%). Several experimental factors differ between the two studies. Although ZH501 and ZH548 RVFV strains have been isolated in the same hospital during the 1977 Egyptian outbreak, they have distinct passaging history [9] and exhibit a small percentage of nucleotide differences [7, 21, 23]. Therefore, we cannot exclude that ZH501 and ZH548 RVFV strains induce distinct survival rates at day 6 p.i. Based on our previous experiments (S1 Table), a lower inoculation dose (10^2^ instead of 10^3^ PFU) is unlikely to be solely responsible for a reduced death rate between days 3 and 6 p.i. in our study. Finally, this difference could be due to mouse sex and genetic background since we used males of the BALB/cByJ inbred strain while the other study was performed on female BALB/c mice, without indication of the substrain. Significant differences between BALB/c substrains have been previously reported with other infectious diseases and immune responses [24, 25], emphasizing the importance of accurately specifying the animal strain used is such studies. Whatever the reason for this difference in survival rates, our findings are consistent with the biphasic RVF disease reported by Smith and colleagues [9] that consists of an acute liver disease followed by a panencephalitis.

Under our conditions, C.MBT-*Rvfs2* mice were highly susceptible to, and died from, the early-onset liver disease, while BALB/c mice overcame it and died later of encephalitis. Our results suggest that one of the mechanisms underlying the increased susceptibility of C.MBT-*Rvfs2* mice is a higher viral replication rate at the cellular level, as shown in primary cultured hepatocytes and by the increased viral load in the liver, despite similar percentage of infected liver cells as assessed by non-quantitative IHC. Altogether, these findings establish the feasibility and exemplify the value of segregating important sub-phenotypes by transferring a single locus, *Rvfs2*, from the early susceptible to a late susceptible background. Susceptibility to liver disease has also been reported in WF inbred rats after subcutaneous infection with RVFV ZH501 [8]. WF rats died by day 2 post inoculation of liver necrosis, whereas LEW rats were resistant to the liver disease but fairly susceptible to the encephalitis [8, 13]. This susceptibility to liver necrosis occurred in a similar time frame after respiratory infection in WF rats [12]. The pathogenic mechanisms that trigger the susceptibility of WF rats and C.MBT-*Rvfs2* mice to RVF hepatic disease may be similar. Indeed, it has been reported that susceptible WF rats had much higher blood viral titers than resistant LEW rats at day 2 p.i. [13], in line with the higher viral production in hepatocytes from WF rats compared with LEW rats [26]. These data suggest that the rat susceptibility locus also controls the production of RVFV. Recently, a major gene for the susceptibility has been mapped within a region on rat Chr 3 [14]. This rat region has homology with mouse Chr 2, indicating that the rat susceptibility locus and *Rvfs2* which maps on mouse Chr 11 do not point at the same gene(s). Therefore, the genetic variations captured in WF rats and MBT mice are different, which makes both rodent models equally interesting and important.

In principle, the RVFV-infected host can protect itself from lethal liver disease using two non-mutually exclusive strategies: (i) resistance to reduce viral burden, (ii) tolerance to reduce the negative impact of the viral burden on host fitness [27, 28]. Our study indicates that the main contributor to early lethality of C.MBT-*Rvfs2* mice can be attributed to higher levels of RVFV in the blood and liver and higher viral replication rate in hepatocytes compared to BALB/c mice. Whether higher susceptibility to liver disease is only due to lower resistance or also to reduced tolerance remains to be determined. The distinct overall phenotype of the BALB/c → BALB/c chimeras compared with the BALB/c → C.MBT-*Rvfs2* and C.MBT-*Rvfs2* → C.MBT-*Rvfs2* chimeras demonstrated an absolute requirement of both BALB/c hematopoietic and BALB/c non-hematopoietic cells for *Rvfs2*’-induced resistance. Previous work, including ours, have revealed the importance of specific genetic regions in the outcome of RVFV infection. This report provides insight into the role of the *Rvfs2* locus. Host genetic factors may influence multiple pathways or molecular or cellular processes, such as the production of virus by infected cells or the immune responses. Our previous work has unraveled an impaired innate immune response in MBT mice [16] as well as other immunological differences [17]. Other studies have emphasized the role of cytokines and T-cell responses in the pathogenesis of RVF disease [29, 30]. The causal variants within the 17 Mb *Rvfs2* genomic interval remain to be identified, as well as the mechanisms by which they control the production of infectious virus particles and the ability to survive the RVF-induced liver disease.

## Materials and Methods

### Ethics statement

Experiments on mice were conducted according to the French and European regulations on care and protection of laboratory animals (EC Directive 2010/63/UE and French Law 2013-118 issued on February 1, 2013). All experimental protocols were approved by the Institut Pasteur Ethics Committee (under #2013-0127, 2016-0013 and dap160063) and authorized by the French Ministry of Research (under #02301, 06463 and 14646, respectively).

### Mice

C.MBT-(JAX0003137-UNC20541010) congenic mice, designated herein as C.MBT-*Rvfs2*, carry a segment of Chr 11 from the MBT/Pas (MBT) inbred strain extending between positions 104,823,629 and 121,738,604 in assembly mapping GRCm37, onto a BALB/cByJ (BALB/c) inbred genetic background [18]. C.MBT-*Rvfs2* and BALB/c mice were bred under specific pathogen-free conditions at the Institut Pasteur.

### Virus production and mouse infection

The RVFV strain ZH548, isolated from a male patient with the acute febrile illness at Zagazig fever hospital, Egypt [21, 31] (obtained from Centre National de Référence des Fièvres Hémorragiques Virales, Institut Pasteur, Lyon, France), was used for all infection studies. All experiments that involved virulent RVFV were performed in the biosafety level 3 (BSL3) facilities of the Institut Pasteur, and carried out in compliance with the recommendations of the Institut Pasteur Biosafety Committee (N° 14.320). Stocks of RVFV ZH548 were titrated by plaque assay on monolayers of Vero E6 cells [32]. Infections were carried out on 9 to 13 weeks old male mice, in BSL-3 isolators. Mice were infected intraperitoneally with 10^2^ PFU of RVFV strain ZH548. Clinical disease scores and mortality were monitored daily for 14 days following infection. Moribund animals were euthanized. Animals that survived were euthanized on the last day of the monitoring period.

### Clinical evaluation

Implantable Programmable Temperature Transponders (IPTT-300) (Bio Medic Data Systems, Inc., Seaford, DEL, USA) were injected subcutaneously into mice one week prior to challenge with RVFV ZH548, and body temperature was monitored daily. Body weight of ZH548-infected mice was measured daily throughout the course of the experiment to evaluate the daily weight loss. Alanine aminotransferase (ALT) and aspartate aminotransferase (ALT) levels were measured using IDEXX diagnostic panels analyzed on a VetTest chemistry analyzer (IDEXX laboratories, Westbrook, ME, USA) on ZH548-infected mice and uninfected controls.

### Viral titer, viral RNA load, and expression of N and NSs viral proteins

Groups of infected BALB/c and C.MBT-*Rvfs2* mice were euthanized on day 3 p.i. Blood was collected by cardiac puncture. The left lateral lobe of the liver was harvested after perfusion from the portal to the cava vein with saline to remove blood-associated RVFV from the tissues. Infectious titers were measured in sera samples and liver homogenates by plaque assay on monolayers of Vero E6 cells [32].

The expression of N and NSs viral proteins was studied by Western blot analysis. Total proteins were extracted from liver samples of two mice used above for viral titration (noted b1, b2, r1 and r2 on figure 6). Protein quantification was done using Micro BCA Protein Assay kit (ThermoFisher Scientific, Waltham, MA). Ten μg of total protein from a cell lysate from AML12 hepatocytes infected with RVFV at an MOI of 3 were used as a positive control. Forty micrograms of total proteins extracted from liver samples and resuspended in Laemmli buffer were run on 14% SDS-polyacrylamide gel and transferred onto nitrocellulose membranes (Amersham, Velizy-Villacoulay, France). Membranes were blocked with a solution of 5% milk (low fat) in PBS containing 0.05% of Tween 20 also used to dilute antibodies (Ab). Proteins were detected by using a rabbit polyclonal Ab raised against a recombinant N protein produced in the baculovirus system, a mouse polyclonal Ab raised against the entire NSs protein [33, 34], or a monoclonal anti-β-actin antibody (A5441, Sigma-Aldrich, Saint-Quentin Fallavier, France). The membranes were incubated with anti-rabbit or anti-mouse Ab coupled to horseradish peroxidase (Sigma-Aldrich,) then reacted with a chemiluminescent substrate (SuperSignal West Dura Extended Duration Substrate, Thermo Scientific), and revealed with G:BOX Chemi chemiluminescence imaging system (Bangalore, India).

### Histology and immunostaining

Groups of infected BALB/c and C.MBT-*Rvfs2* mice were euthanized at different times along the 14-day period of observation to monitor the development of RVF disease. A first group was euthanized at an early stage of infection, day 3 p.i. A second group was euthanized at the first clinical signs of illness which occurred on day 3 or 4 p.i. in C.MBT-*Rvfs2* mice, and between days 6 and 9 p.i. in BALB/c mice. Finally, BALB/c mice that survived until day 14 p.i. were also euthanized. Non infected BALB/c and C.MBT-*Rvfs2* mice were used as controls. The liver and brain were removed and immediately fixed for one week in 10% neutral-buffered formalin for biosafety reasons. Samples from each organ were embedded in paraffin; 4 μm-thick sections were cut and stained with hematoxylin and eosin (HE). Microscope slides were coded for blinded studies, and examined by a qualified veterinary pathologist (GJ). Non-quantitative immunohistochemical detection of the RVFV-infected cells was done using mouse antibodies against the N protein (dilution 1:100) [35]. A rabbit monoclonal antibody (Ref: AB16667, dilution 1:50; Abcam, Paris, France) was used to detect Ki67 antigen. Visualization was performed with the Histofine Simple Stain MAX-PO kit (Nichorei Biosciences Inc., Tokyo, Japan), a labeled polymer prepared by combining amino acid polymers with peroxidase and secondary antibody which is reduced to Fab’ fragment. This visualization procedure allows amplification of the positive signal and limitation of the background staining, especially when using mouse antibodies as for the detection of the RVFV. However, it does not allow quantitative evaluation of positive signals and intensity comparisons between samples.

### Primary hepatocyte preparation and infection

Seven to 12 week-old BALB/c and C.MBT-*Rvfs2* male mice were euthanized by cervical dislocation. Suspensions of hepatocytes were prepared as described in Li et al. [36] using Collagenase type IV (PAN Biotech, Worthington, UK) at 100U/ml. Cells were cultured also according to Li et al. [36]. On the day after preparation, hepatocytes were infected with RVFV at an MOI of 3 for 1 hr. At 15, 24, 48 and 60 hours post-infection, the supernatant was collected for virus titration by plaque assay as above. Each condition was done in triplicate (3 wells).

### Bone marrow transplantation

Bone marrow cells (BMCs) were collected from both tibias and femurs of 5-6 week-old BALB/c or C.MBT-*Rvfs2* donor male mice. BMCs were resuspended in Hanks’Balanced Salt Solution. After irradiation with one sub-lethal dose of gamma radiation (700 rad), 5-6 week-old BALB/c or C.MBT-*Rvfs2* recipient male mice received ~ 3×10^6^ BMCs in 0.15 ml by intravenous injection in the retro-orbital sinus. The extent of reconstitution was evaluated using a semi-quantitative PCR assay based on primers to *Apoptosis-associated tyrosine kinase (Aatk)* gene (Forward, 5’-CTACCCCAGGAGGACTGTGTCAGG-3’ and reverse 5’-GTCCTCCCCAACAATATCCTGGTGC-3’) that maps within *Rvfs2* interval. BALB/c and MBT alleles produce a fragment of 180 bp and 127 bp, respectively. Six weeks after the transplantation, the reconstitution in total peripheral blood of (BALB/c → C.MBT-*Rvfs2*) and (C.MBT-*Rvfs2* → BALB/c) mice was higher than 90%. At that time, bone marrow chimeras were infected intraperitoneally with 100 PFU of RVFV strain ZH548.

### Statistical analysis

Statistical analysis was performed using GraphPad Prism 6.0 (GraphPad Software, La Jolla, CA, USA) and R softwares. Mantel-Cox’s Logrank test was applied to assess survival curve differences. Two-way ANOVA was used to assess body weight and body temperature differences, with the two factors being the strains and the days post-infection. The p-values shown indicate the significance of the difference between strains. Mann Whitney-U test was used to analyze viral titers in serum and liver.

## Acknowledgments

The authors are grateful to Daniela Faust for valuable advices in primary hepatocyte preparation. They thank Jean Jaubert and Rashida Lathan for their encouragement and fruitful discussion. The technical assistance of Patricia Flamant, Huot Khun, and Carole Tamietti is gratefully acknowledged. The authors also thank Clémentine Dillard and Marine Perey for technical help during their short stay in the laboratory.

## Data Availability Statement

All relevant data are within the paper.

## Author Contributions

Conceptualization: LB, XM, JJP

Formal analysis: LB, DS, DH, XM, JJP

Funding acquisition: GJ, MF, JJP

Investigation: LB, GJ, DS, DH, OBD, MB, ST, TZDV

Methodology: GJ, AC, MF, XM, JJP

Project administration: JJP

Resources: GJ, MF, XM, JJP

Supervision: XM, JJP

Validation: LB, GJ, MF, AC, XM, JJP

Visualization: LB, GJ, XM, JJP

Writing – original draft: LB, XM, JJP

Writing – review & editing: LB, GJ, DS, DH, MB, ST, TZDV, MF, XM, JJP

## Supporting Information

**S1 Table.** Survival rate and days of death of MBT/Pas and BALB/cByJ 9-12 week-old male mice infected with 10, 100 or 1000 PFU of ZH548 RVFV (dpi : days post-infection).

